# Folate receptor targeted nanoparticles containing niraparib and doxorubicin for treatment of high grade serous ovarian cancer

**DOI:** 10.1101/2022.08.29.505711

**Authors:** Lucy Wang, James C. Evans, Lubabah Ahmed, Christine Allen

## Abstract

Combination chemotherapy is an established approach used to manage toxicities while eliciting an enhanced therapeutic response. Delivery of combinations of drugs in specific molar ratios has been considered a means to achieve synergistic effects resulting in improvements in efficacy while minimizing dose related adverse drug reactions.

The benefits of this approach have been realized with the FDA approval of Vyxeos^®^, the first liposome formulation to deliver a synergistic drug combination leading to improved overall survival against standard of care. In the current study, we demonstrate the synergistic potential of the PARP inhibitor niraparib and doxorubicin for the treatment of ovarian cancer. Through *in vitro* screening in a panel of ovarian cancer cell lines, we find that niraparib and doxorubicin demonstrate consistent synergy/additivity at the majority of evaluated molar ratio combinations.

Further to these findings, we report formulation of a nanoparticle encapsulating our identified synergistic combination. We describe a rational design process to achieve highly stable liposomes that are targeted with folate to folate-receptor-alpha, which is known to be overexpressed on the surface of ovarian cancer cells. With this approach, we aim to achieve targeted delivery of niraparib and doxorubicin at a pre-determined synergistic molar ratio via increased receptor-mediated endocytosis.

**Graphical Abstract:** 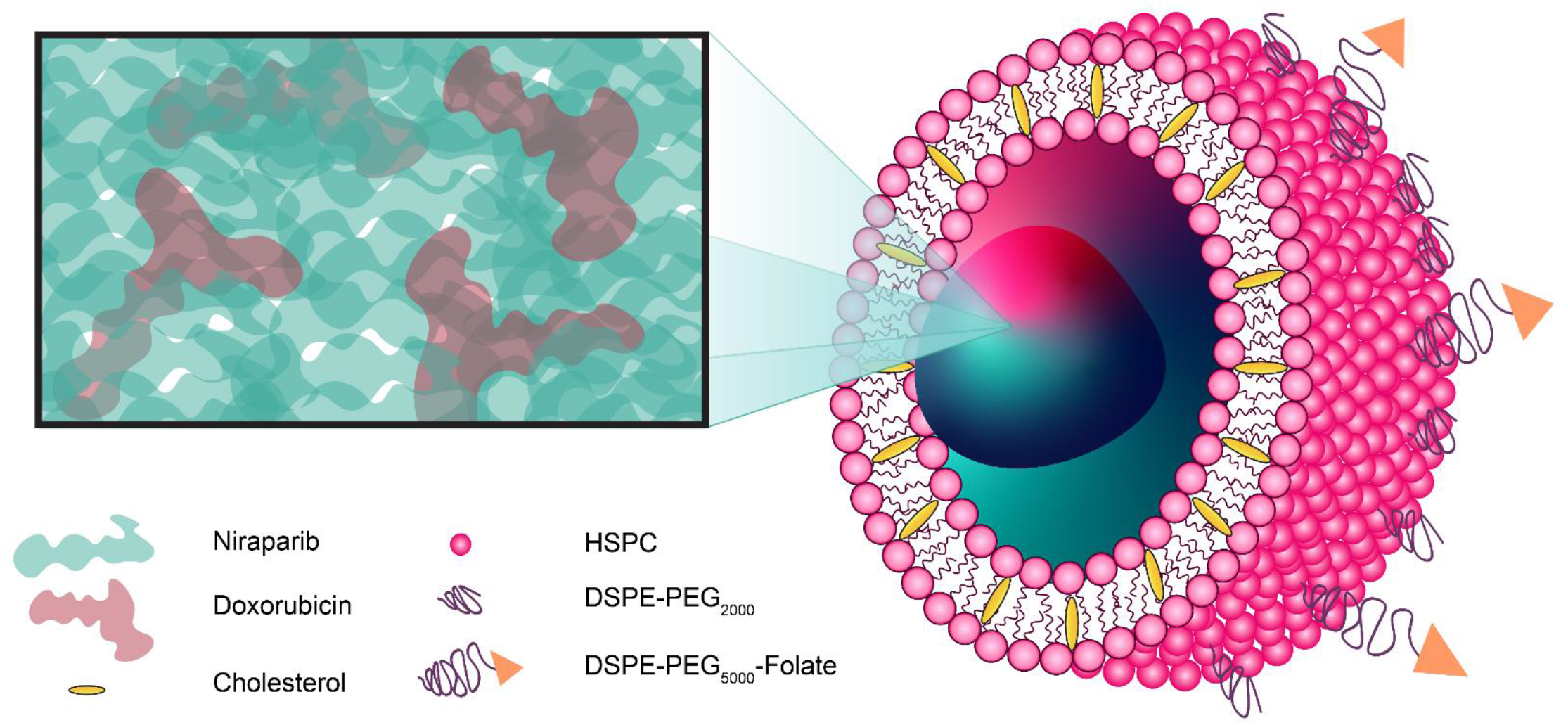

## 1. Introduction

Ovarian cancer (OC) is the fifth leading cause of cancer related deaths among Canadian and American women, with the poorest five-year survival rates among all gynecological cancers [1,2]. Despite significant research in this area, patient survival outcomes have seen relatively small gains in survival rates with average age adjusted rates of death falling by merely 2.7% per year [3].

Nevertheless, within the past decade we have seen the first major innovation in OC treatment since the 1990s with the FDA approval of the first-in-class poly-ADP ribose polymerase (PARP) inhibitor (PARPi), olaparib (OLP). In 2014, OLP was approved as a monotherapy for patients with germline BRCA-mutated advanced stage OC who have undergone prior treatment with three or more chemotherapies (such as taxanes and platinum agents) [4]. OLP use has since been expanded to include maintenance therapy in recurrent OC patients who are either fully or partially responsive to platinum-based therapy [5].

PARPis are a family of molecular therapies that take advantage of the homologous recombination repair defects (i.e. the “BRCAness” phenotype) prevalent in the majority of high grade serous ovarian cancer (HGSOC) tumors to create a synthetic lethality effect [6,7]. There are currently three PARPis approved for the treatment of OC: OLP, rucaparib, and niraparib (NIRB), that differ with respect to their degree of PARP inhibition and PARP trapping [8,9].

To improve patient outcomes, there have been several studies aimed at determining the therapeutic potential of PARPis in combination with other chemotherapeutic and molecular therapy agents. Clinical trials that add PARPis to traditional first- or second-line OC chemotherapies have garnered mixed results with key limitations being dose limiting toxicities. These toxicities have been reported to be mainly hematological in nature and sometimes affect as many as two-thirds of the cohort at even low dosage levels [10–18]. Furthermore, recent work looking at combining PARPis with novel molecular therapies such as ATR and WEE1 inhibitors have found similar hematological toxicity drawbacks [19,20].

Although sequential combination therapy and the use of drug “holidays” have been proposed (and successfully applied) to overcome toxicity barriers, administration of concomitant drug combinations remains ideal for ease of clinical application and potential improved effectiveness that oftentimes cannot be assessed in a head-to-head comparison to sequential therapy.

Herein we propose to reduce PARPi related concomitant drug combination toxicities by encapsulating combination cancer drugs that have undergone *in vitro* pre-screening for synergistic action into nanoparticles for both passive and active delivery to cancer cells. The benefits of nanoparticle mediated combination delivery combine the well documented reduced systemic toxicity effect of nanoparticle encapsulation with the ability to package and deliver optimized synergistic drug combinations at the relative ratio found to be most effective.

This optimized “ratiometric” advantage to applying drug combinations is well documented in the literature [21–24]. Vyxeos^®^ is the first dual-encapsulated nanoparticle that uses this approach to treat acute myeloid leukemia. Delivering a synergistic 5:1 molar ratio of cytarabine and daunorubicin, Vyxeos is the only nanoparticle formulation to date that has demonstrated superior overall survival when compared to the standard of care (i.e. 7-day continuous infusion of cytarabine with 3-day infusion of daunorubicin) [25,26]. Furthermore, patients who received Vyxeos (as opposed to the standard of care) had significantly lower incidence of cardiac toxicity [27]. These outcomes provide the basis for our current approach to the toxicity associated with PARPis when combined with traditional standard of care chemotherapy.

Using the Chou-Talalay method, we have previously demonstrated that a combination of the PARPi, OLP and anthracycline, doxorubicin (DOX) exhibit synergy at a range of specific relative molar ratios in a panel of nine OC cell lines [28]. It is thus hypothesized that NIRB and DOX could also have similar potential for synergy. In the following we describe the *in vitro* screening of various molar ratios of the PARPi, NIRB and DOX in a panel of OC cell lines. Subsequently, we outline the rational design of an actively targeted, highly stable, dual-encapsulated NIRB and DOX liposome for the treatment of HGSOC.

## 2. Results

### 2.1 NIRB and DOX drug combination screening

To elucidate the synergistic potential of our test compounds, NIRB and DOX were combined at varying relative molar ratios and screened for cytotoxicity in a panel of representative HGSOC cell lines. The combination index (CI) values were determined using the CompuSyn software. Thresholds for synergism, additivity, and antagonism were set at ≤0.9, between 0.9 and 1.1, and ≥1.1, respectively.

As shown in Figure 1a-b, CI values obtained for ratiometrically combined NIRB and DOX combinations applied to cell monolayers showed a general trend of mean additivity for all applied molar ratios at both fractions affected (Fa) 0.5 and 0.75 (i.e., 50% and 75% cell death, respectively). Interestingly, pockets of mean antagonism were exclusively observed in the PEO4 cell line, a platinum resistant sister pair to the PEO1 cell line.

**Figure 1:**
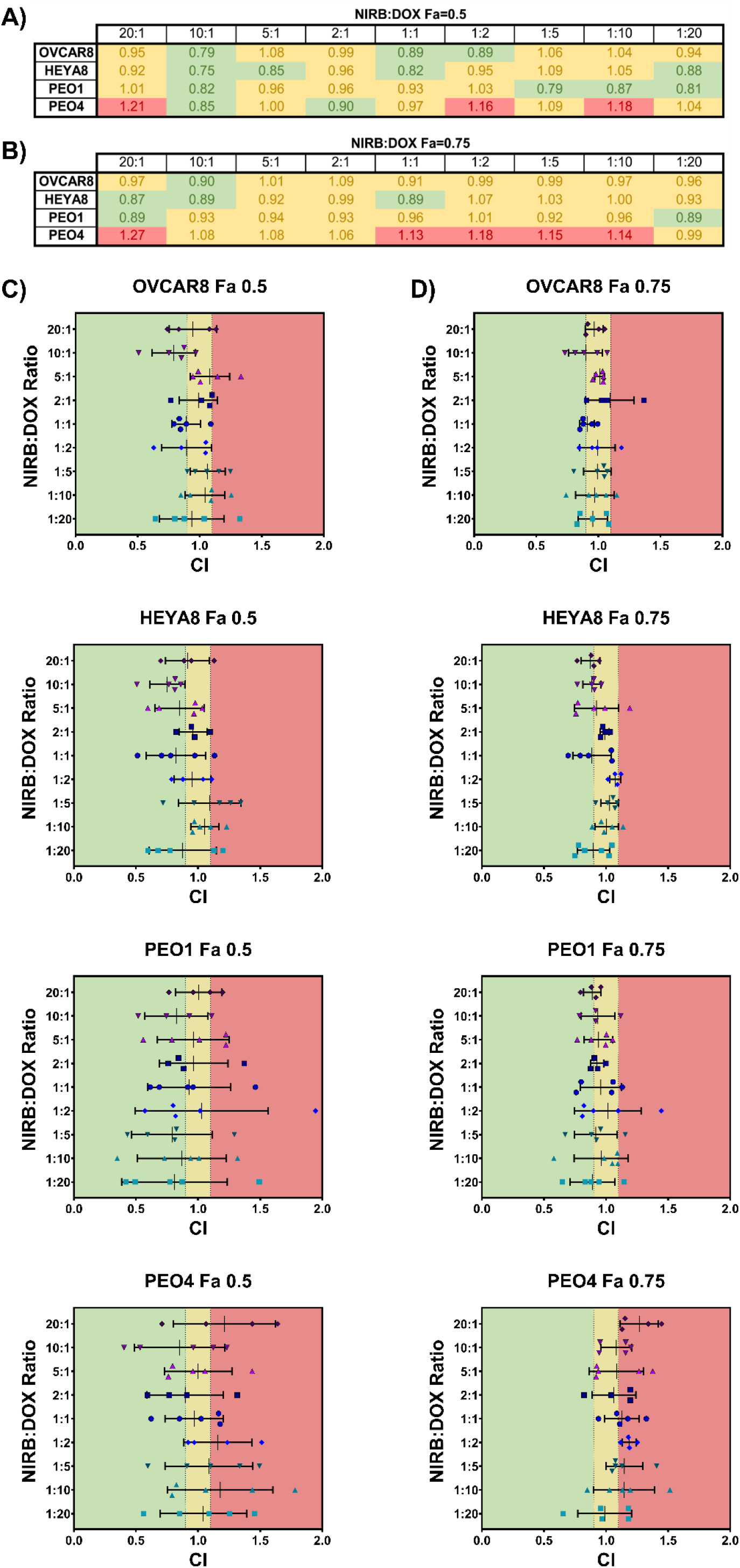
Heat map and scatter plot showing either A-B) mean or C-D) mean±SD CI values for treatments of NIRB:DOX at varying relative molar ratios on a panel of four HGSOC cell lines. CI values ≤0.9 indicates synergistic effects (green), 0.9>CI<1.1 indicates additive effects (yellow), and CI≥1.1 indicates antagonistic effects (red). The 10:1 molar ratio is shown to have consistent synergistic/additive effects in all cell lines at both Fa=0.5 and Fa=0.75 except for PEO4, wherein the combination of NIRB and DOX seems to exhibit a wide variation including antagonism across all molar drug ratios. CI values were calculated using CompuSyn software.

As such, the 10:1 NIRB:DOX ratio was the only test mixture that showed consistent mean synergy across all tested cell lines at Fa=0.5 with only a slight increase towards mean additivity at Fa=0.75.

Expressed as scatter plot, Figure 1c-d shows that the standard deviations of biological replicates for the 10:1 ratio consistently remain in the synergistic and additive range for all cell lines except PEO4. PEO4 was also found to be the only cell line where variability for all tested ratiometric mixes deviated into the antagonistic region, a detail that is not conveyed when solely looking at mean values.

### 2.2 Cell surface expression of folate receptor alpha (FRα) and cellular uptake of folate conjugated liposomes

Given that 70% of primary and 80% of recurrent HGSOC tumors overexpress FRα, the folic acid small molecule was identified as a potential active targeting ligand [29]. Per Figure 2a, surface expression of FRα on the selected panel of OC cell lines was measured by flow cytometry with the breast cancer cell line MCF7 used as a negative control. The OVCAR8 cell line was found to have significantly higher FRα expression compared to both the negative control, HEYA8 and PEO1 cell lines. OVCAR8 was thus selected to evaluate the suitability of our carrier for *in vitro* cellular uptake.

**Figure 2:**
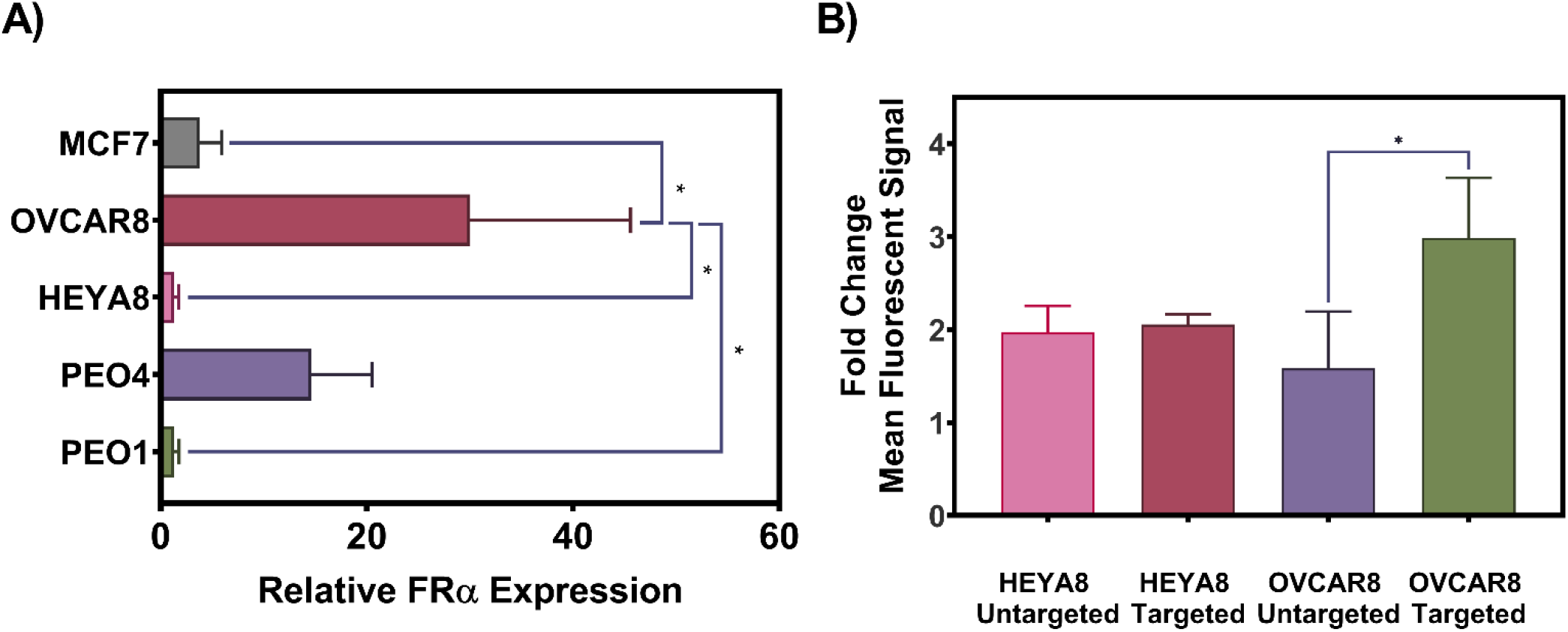
A) Relative FRα expression in the selected panel of OC cell lines with MCF7 (breast cancer cell line) as a negative control and B) Cellular uptake of folate conjugated versus unconjugated fluorescent calcein loaded liposomes. Flow cytometry was used to confirm the relative expression of FRα in the panel of OC cell lines. Data is presented as mean FITC for treated cells divided by the mean FITC for untreated cells, * p<0.05 (n=3). Flow cytometry was used to determine the uptake of folate conjugated and folate unconjugated calcein-loaded liposomes in FRα high (OVCAR8) and low (HEYA8) expressing cells, presented as mean FITC values. Data is presented as mean FITC for treated divided by the mean FITC for untreated cells, * p<0.05 (n=3).

As shown in Figure 2b, cellular uptake of folate conjugated, fluorescent probe encapsulating liposomes into OVCAR8 cells was significantly greater than unconjugated control liposomes. This effect was not observed in the HEYA8 cell line which does not overexpress surface FRα.

### 2.3 NIRB and DOX dual loading optimization

Dual loading of NIRB and DOX into a common liposome was pursued to deliver a controlled, synergistic ratio of both drugs to cancer cells. Remote drug loading using a pH gradient and triethylammonium sucrose octasulfate (TEA_8_SOS) as an internal entrapment buffer was selected as the method of choice. NIRB and DOX were added to blank liposomes at varying relative molar ratios to study the dual loading pattern and to determine if one drug loads preferentially into liposomes in comparison to the other. Successfully entrapped NIRB:DOX molar ratios were plotted as a function of added NIRB:DOX molar ratios to elucidate a potential relationship (Figure 3).

**Figure 3:**
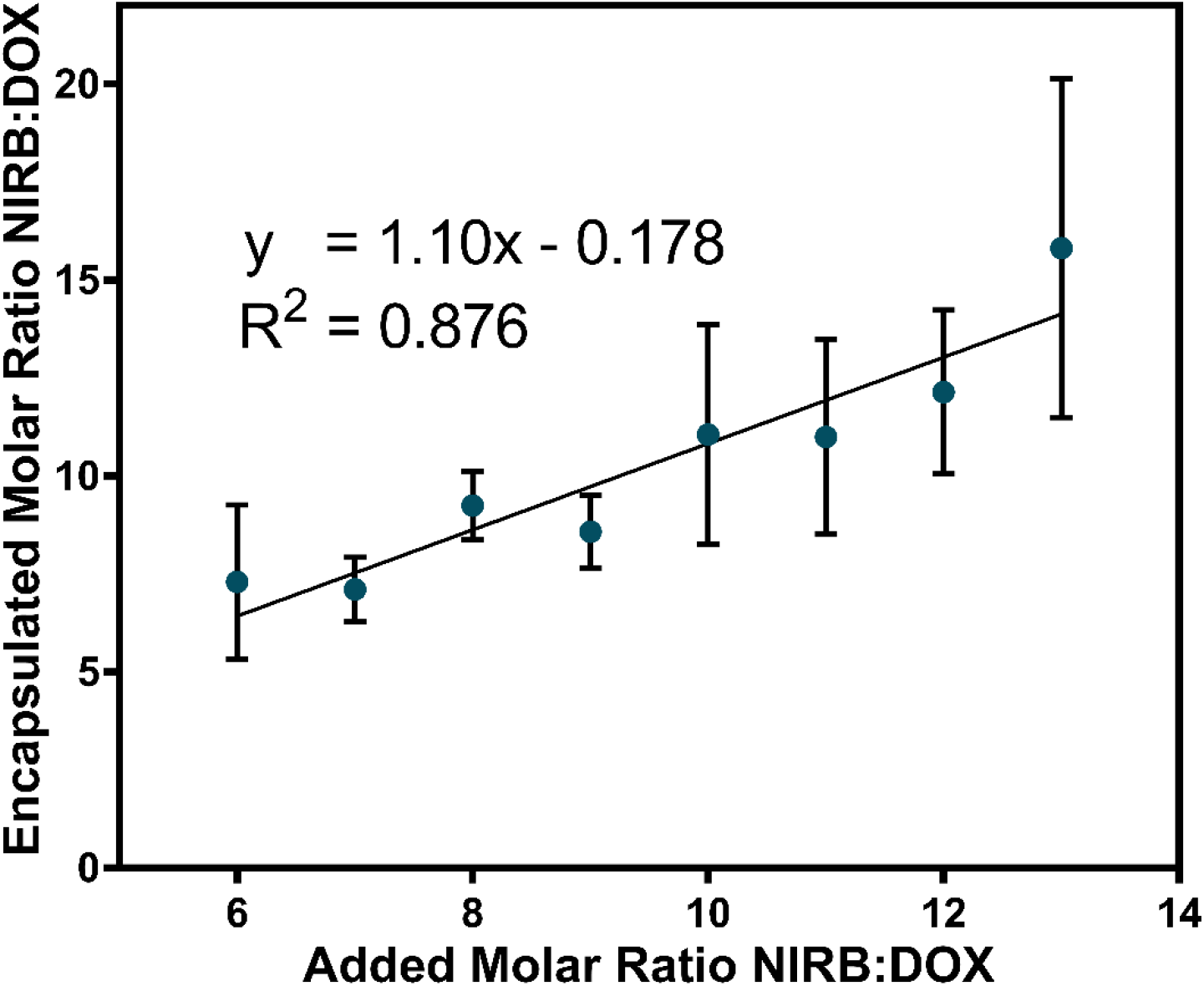
Relationship between the added molar ratio of NIRB:DOX and the corresponding resulting “encapsulated molar ratio” following active loading with TEA_8_SOS as a trapping agent. DOX and NIRB concentrations in the resultant liposomes were quantified by mass spectrometry. Increases in relative molar ratios of encapsulated NIRB:DOX is related to increases in relative molar ratios of added NIRB:DOX with an R^2^=0.876. The determination of this relationship allowed for the formulation of a synergistic NIRB:DOX liposome in a predictable and reproducible fashion. Data is presented as mean±SD. (n=3).

NIRB and DOX were found to dually load into blank liposomes, containing TEA_8_SOS in their internal aqueous core, in a linear relationship wherein increasing the added proportion of NIRB relative to DOX at the active loading step increased the encapsulated amount of NIRB relative to DOX in a predictable linear manner. This linear relationship can be expressed as *y* = 1.10*x* – 0.178 between the tested limits with an *R*^2^ = 0.876.

Substituting *y* = 10, the predicted ratio of added NIRB:DOX was determined to be 9.24:1. Applying this experimentally, the resultant folate conjugated liposomes (NIRB-DOX-FA) reproducibly encapsulated 9.4±0.6:1 molecules of NIRB relative to DOX. Table 1 shows a summary of the characteristics of the folate conjugated and drug loaded liposomes.

**Table 1:** Size, encapsulation efficiency, and drug to lipid ratio of the optimized, folate conjugated, lead liposome candidate, NIRB-DOX-FA. Nanoparticle size was determined by DLS. NIRB and DOX concentrations were determined by HPLC-UV and lipid quantification was determined by HPLC-ELSD.

| Size (nm) | Encapsulation Efficiency (%) |  | Drug:Lipid Ratio | Resultant NIRB:DOX Molar Ratio |
| --- | --- | --- | --- | --- |
|  | NIRB | DOX |  |  |
| 124.6±53.9 | 87.6±19.9 | 71.5±17.3 | 0.4±0.1 | 9.4±0.6 |

**Table 2:** Selected panel of human OC cell lines and their relevant mutations, culture conditions, and 96-well plate seeding densities used for *in vitro* combination cytotoxicity studies.

| Cell Line | Characteristic Mutations | Culture Media | Seeding Density |
| --- | --- | --- | --- |
| OVCAR8 | TP53 [49], BRCA1 hypermethylation [50] | RPMI, 10% FBS, 100 units/mL penicillin and 100 μg/mL streptomycin | 1000 cells/well |
| HEYA8 | PIK3CA, KRAS, BRAF [51] |  | 1000 cells/well |
| PEO1 | TP53, BRCA2 [31] |  | 3000 cells/well |
| PEO4 | TP53 [31] |  | 3000 cells/well |

### 2.4 NIRB-DOX-FA liposome *in vitro* characterization

The *in vitro* release of drug from the NIRB-DOX-FA liposomes was evaluated under sink conditions that mimic blood serum (Figure 4a). Timepoints collected were 0-hours (i.e., sample run down the size exclusion column immediately after addition of release media to purified liposomes), 1-hour, 2-hours, 4-hours, 6-hours, 10-hours, 24-hours, 48-hours, 72-hours, 96-hours, and 120-hours.

**Figure 4:**
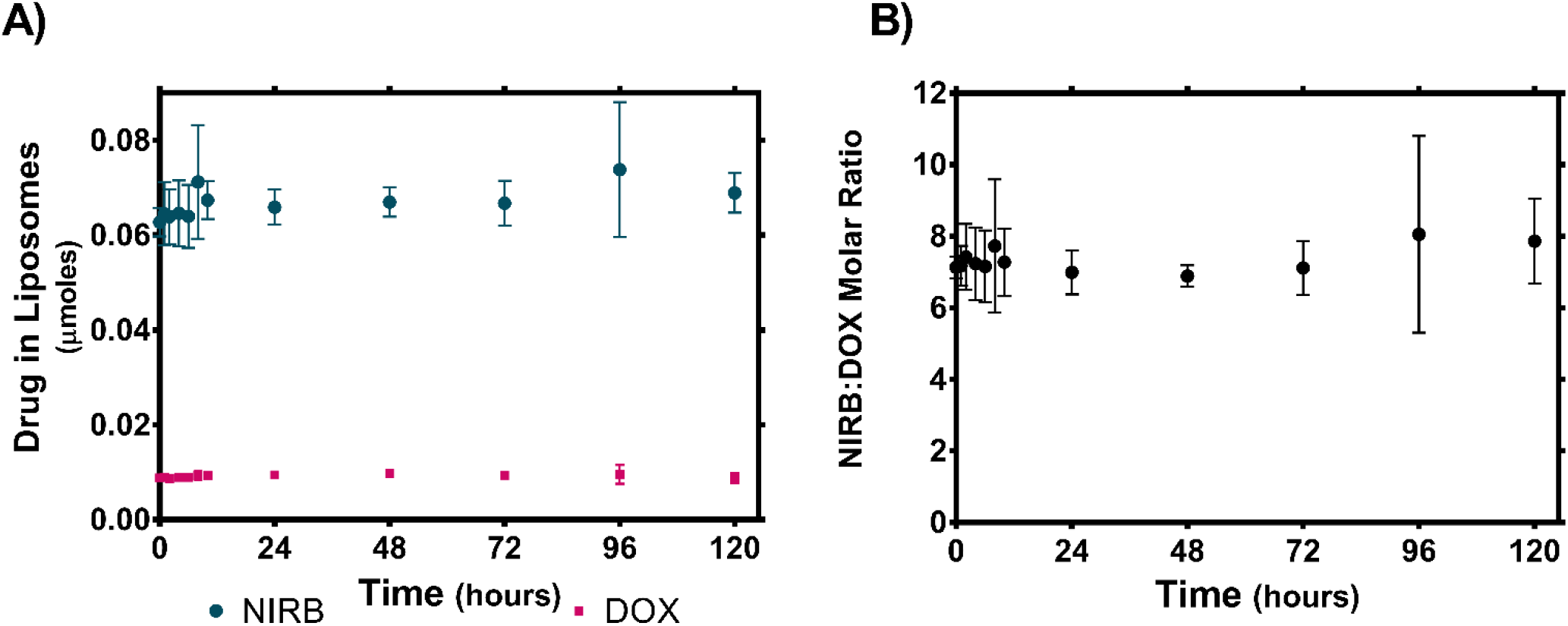
A) *In vitro* release of NIRB and DOX from NIRB-DOX-FA in BSA and the B) corresponding encapsulated NIRB:DOX molar ratio. The y-axis shows the mean absolute value (μmoles) of NIRB and DOX remaining in the liposomes (±SD) within collected 100μL fractions when NIRB-DOX-FA liposomes are subjected to biologically relevant (i.e., BSA; 50 mg/mL) sink conditions over five days. Virtually no drug was released between time 0-hours and 120-hours, resulting in the maintenance of initial versus final NIRB:DOX molar ratios (7.13±0.30:1 at time 0-hours and 7.86±1.19:1 at time 120-hours). Data is presented as mean±SD. (n=3).

As shown in Figure 4b, the NIRB:DOX molar ratio was maintained within 7.3±1.1 throughout the entire 5-day study period. This observed NIRB:DOX ratio deviates from the target 10:1 synergistic ratio but remains within the synergistic/additive range for all cell lines.

It is hypothesized that this discrepancy is a result of bead adherence and drug loss on the size exclusion column upon purification of release samples – as evidenced by the lower than anticipated time 0-hours NIRB:DOX molar ratio (7.1±0.3). When all timepoints are compared to 0-hours, there were no statistically significant changes in encapsulated ratio. As such, relative to timepoint 0-hours, it is concluded that there was no measurable release of drug payload over a 5-day incubation period.

Cryo-transmission electron microscopy of unloaded vs loaded liposomes were conducted to elucidate the particle morphology of the dual drug loaded liposomes. Representative images of the drug loaded liposomes reveal dark areas of high electron density that are not present in unloaded liposomes (Figure 5). Using ImageJ software to analyze the mean gray area difference across the phospholipid membrane of loaded vs unloaded liposomes reveals that drug loading results in a significant increase in mean gray area difference across the liposomal membrane (Figure 6). In addition, drug loading and entrapment using the TEA_8_SOS agent is shown to change the liposomal morphology from a circular to an oblong shape.

**Figure 5:**
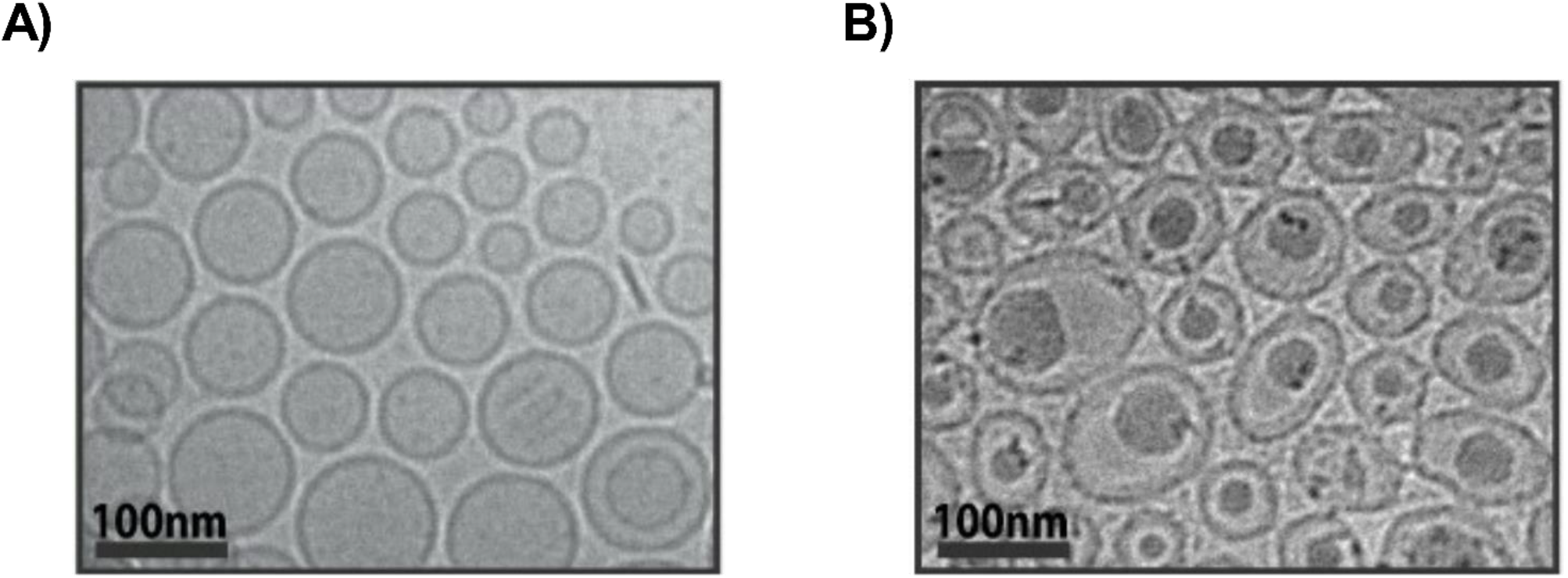
Cryo-TEM images of A) unloaded TEA_8_SOS liposomes and B) NIRB and DOX loaded TEA_8_SOS liposomes. Darker versus lighter areas demonstrate the presence of a greater versus lower electron density, respectively. Comparing images of loaded versus unloaded liposomes, drug loaded liposomes exhibit dark, electron dense deposits at the core. These deposits may be due to precipitated drug in the liposomal core.

**Figure 6:**
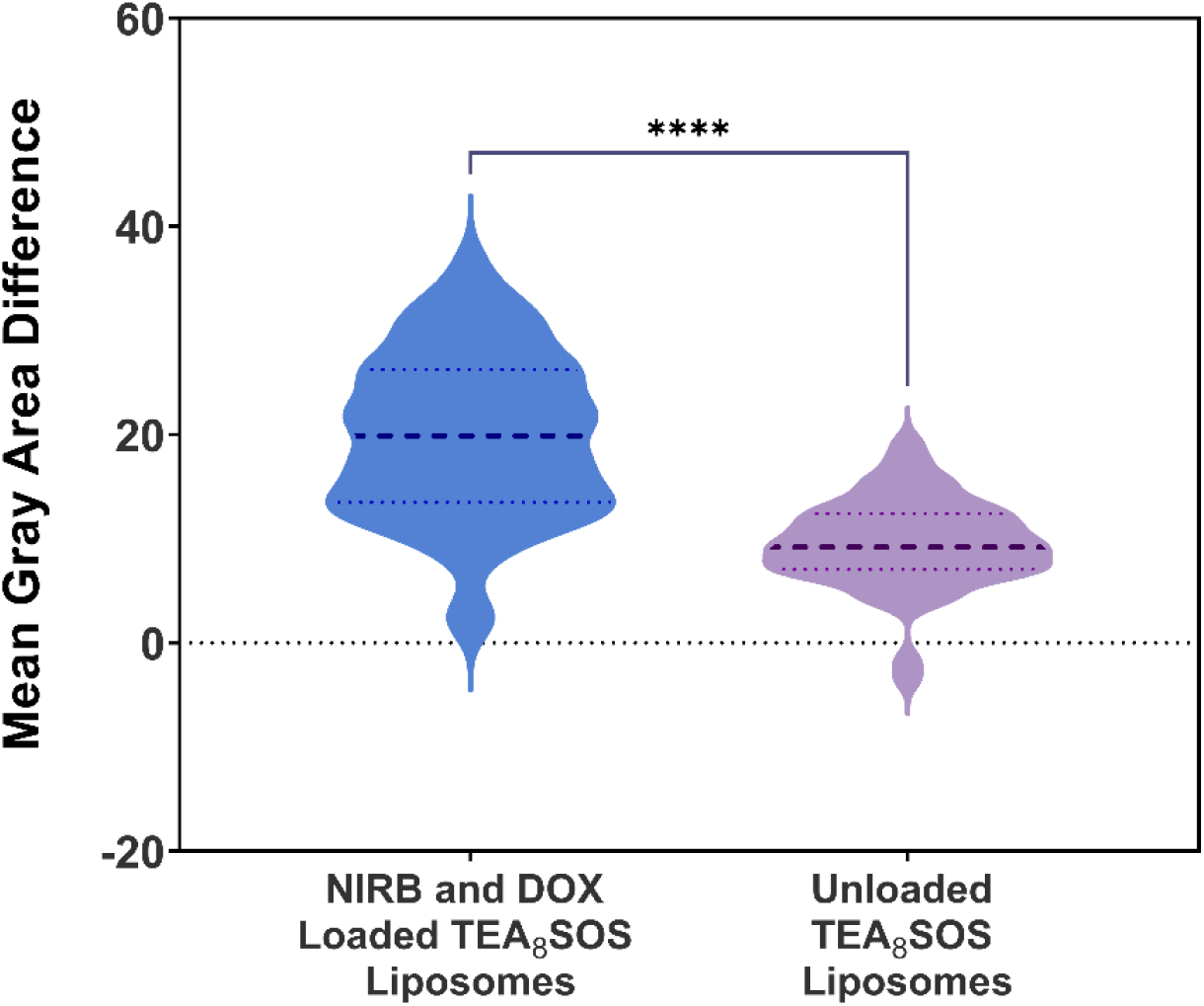
Differences in mean gray area of the exterior versus interior of drug loaded (NIRB-DOX-FA) and unloaded TEA_8_SOS liposomes. Mean gray area is defined as the average photo intensity divided by the number of pixels in a selected image area. Mean gray area was determined to quantify the darkened electron dense areas at the core of NIRB-DOX-FA liposomes versus unloaded liposomes. As such, statistically significant differences in mean gray area across the bilayer of NIRB-DOX-FA liposomes, relative to unloaded liposomes, highlights the presence of electron dense drug deposits after active drug loading. Mean gray area of the liposomes was determined by ImageJ software for the cryo-TEM images. Data is presented as differences in mean gray area (n=56 liposomes, **** p<0.0001).

## 3. Discussion

### 3.1 Synergy in the NIRB and DOX combination is both ratiometrically and homologous recombination repair proficiency dependant

The NIRB:DOX combination demonstrated consistent mean additivity with regions of synergy for most molar ratios tested across all cell lines with the exception of the PEO4 cell line (Figure 1a-b). Among the screened molar drug ratios, the 10:1 molar ratio of NIRB:DOX demonstrated the most consistent synergistic effect across most cell lines, highlighting 10:1 as a combination candidate for nanoparticle delivery. Synergy of the 10:1 ratio remains true when looking at CI values in terms of mean±SD (Figure 1c-d). While the variability of CI values for most applied NIRB:DOX molar ratios include antagonism, the CI standard deviation of the 10:1 ratio consistently falls within the synergistic/additive range for all cell lines except for PEO4.

Though the mechanism by which the NIRB and DOX combination results in either synergy or antagonism is not fully understood, it is hypothesized that the combination of DOX and NIRB could exhibit synergy via the creation of double stranded DNA (dsDNA) breaks. Previous work exploring the synergistic mechanisms of DOX when combined with another PARPi, OLP, revealed that the PARPi-anthracycline combination exhibits its synergistic cytotoxicity by creating more dsDNA breaks than either drug applied as a monotherapy [28]. Though comparative mechanistic studies have not been conducted with NIRB and DOX, given the PARP trapping and inhibition similarities between OLP and NIRB, it is possible that the NIRB and DOX combination exhibits synergy via similar mechanisms [30].

Furthermore, closer examination of the synergistic potential of the NIRB:DOX combination when applied to PEO4 reveals a general trend towards antagonism at all tested molar ratios. These results were consistent with previous work done in our group which showed a similar antagonistic pattern with the OLP and DOX combination across all tested molar ratios [28]. These findings are in stark contrast to the synergistic/additive profile of the NIRB and DOX combination shown in the PEO1 cell line.

This comparison is critical to our understanding of PARPi and DOX combinations as PEO1 and PEO4 are sister cell lines that were derived from the same patient at different points of disease progression. The PEO1 cell line was obtained from the patient at the stage of primary tumor debulking following one round of neoadjuvant chemotherapy (cisplatin, 5-FU, and chlorambucil) and, PEO4 was removed during the secondary tumor debulking after the development of clinical resistance to the aforementioned chemotherapy [31]. Inspection of the PEO1 and PEO4 sister pair reveals a BRCA2 mutation reversion to wild-type that is present in the PEO4 cell line [32]. Given that BRCA2 works exclusively on the homologous recombination (HR) repair pathway, we hypothesize that the observed synergy for the NIRB and DOX combination is highly dependent on HR repair deficiency [33]. However, additional studies are needed to support this hypothesis.

These results suggest that a synergistic effect for the combination of NIRB and DOX is both cell line and molar ratio dependent. Translation of this combination to the clinic would thus likely need to be in conjunction with criteria for identifying patients likely to respond to this therapy. In the case of OLP, during AstraZeneca’s 2014 Phase I trial, the oral PARPi OLP and intravenous (IV) pegylated liposomal doxorubicin (PLD) combination elicited either a complete or partial response in only 25% of platinum resistant patients as opposed to the observed 71% for platinum sensitive patients [34]. The present study aimed to investigate whether a ratiometric approach to PARPi and DOX combinations would yield drug synergy, potentially bridging the difference in observed response rate between platinum sensitive and resistant disease. Based on the limited number of cell lines evaluated, HR repair proficient tumors may be an exclusion criterion for this treatment strategy and may be useful to identify good vs bad responders. However, this warrants further investigation.

Clinical trials conducted on oral NIRB (Zejula^®^) and IV PLD in patients with platinum resistant/refractory HGSOC have been attempted. It was hypothesized that the combination of Zejula and PLD would result in either a *“tumor response rate equal or superior to that of historical data for [PLD] alone”* or a reduction in *“number of participants with dose-limiting toxicities”* (NCT01227941). This study has since been terminated with no published data of the findings.

Though it is not known why this specific trial was terminated, it is possible that the platinum resistant/refractory HGSOC cohort contained patients with HR proficient tumors which, based on our findings, could be less sensitive to the synergistic action of PARPi and DOX. The heterogenous HR proficiency background of patients in the refractory disease setting would also be more likely given that BRCA reversion is a known mechanism of treatment resistant OC, particularly after long term use of PARPi in the maintenance setting [35].

All this points to our need for a better understanding of how next generation molecular therapies affect cancer cell biology and their management of intervention induced genomic instability. Given that the HR repair pathway and its relation to the larger, overall DNA damage response (DDR) network is vastly complex, our work to capitalize on innate cancer cell deficiencies in these pathways to create synthetic lethality responses with not one, but a cocktail of modern-day therapeutics must be further mechanistically studied to better inform decisions that are made in the clinic.

### 3.3 Active targeting ensures cellular uptake of synergistic payload

One of the marked issues with applying combination drug therapies at specific optimized ratios is delivery. While physiological challenges consist of differing pharmacokinetic and biodistribution properties of each individual compound (both separately and when administered together), the cellular challenge consists of controlling for and ensuring cellular uptake at the suggested ratio. For both barriers to implementation, targeted delivery of a dual loaded nanoparticle provides a feasible solution. While encapsulation of optimized drug combinations regulates the PKBD profile of the drug cocktail to a singular profile, active targeting for cellular uptake ensures controlled cellular uptake of the optimized ratio.

As such, active targeting using the folic acid small molecule was pursued. As mentioned previously, the cell surface FRα receptor is overexpressed in the majority of both primary and recurrent HGSOC tumors. The selection of FRα as the target of choice was informed by literature demonstrating enhanced efficacy of folate receptor targeted formulations in preclinical animal models relative to their untargeted controls [36–38]. In the clinical setting, novel therapeutics targeting FRα have seen success in the clinic. A notable example in this category include mirvetuximab/IMGN853, an antibody drug conjugate that led to significant improvements in secondary outcomes during phase III clinical trials [39].

To assess whether folate targeting can facilitate improved cellular uptake of dual-loaded liposomes into FRα overexpressing cells, the expression status of FRα in the OC cell panel was determined to identify an appropriate FRα overexpressing model. As shown in Figure 2a, the OVCAR8 cell line exhibited significantly higher levels of FRα expression relative to the MCF7 negative control (p<0.01). In contrast, none of the other cell lines evaluated (i.e., HEYA8, PEO1/4) demonstrated enhanced expression of FRα relative to MCF7.

Given the FRα expression profiles, cellular uptake of folate conjugated versus unconjugated liposomes was then evaluated in the FRα overexpressing OVCAR8 cells versus the FRα non-overexpressing HEYA8 cells. Statistically significant increases in cellular uptake of folate conjugated versus unconjugated liposomes in the OVCAR8 cell line compared to the HEYA8 cell line determined our carrier lipid composition and active folate targeting strategy to be effective in improving liposomal payload uptake into FRα overexpressing cells.

### 3.4 Optimized liposomal dual encapsulation of NIRB:DOX capitalizes on observed synergy

Given that the observed *in vitro* combination results only demonstrated consistent pronounced synergy at a 10:1 molar ratio (NIRB:DOX), it was imperative that this molar ratio be delivered to cancer cells with little deviation. To achieve this, liposomal core composition was optimized to deliver a controlled amount of each drug into OC cells.

As described in a recent review on nanocrystallization by the Boyd group, the physical state of drug at the nanoparticle core can significantly influence its release kinetics. A solid core morphology has been shown to slow the rate of drug release [40]. This phenomenon has been extensively studied for DOX loaded liposomes and the crystal structure of the resulting drug precipitates [41]. It was hence hypothesized that, to achieve minimal systemic drug release, the proposed NIRB-DOX co-encapsulated formulation would benefit from a more solid core morphology.

With regards to ionizable drugs such as NIRB and DOX, it was prudent to explore gradient mediated active loading using entrapment salts such as TEA_8_SOS. Developed by Drummond and colleagues, the TEA_8_SOS entrapment agent has been shown to form stable complexes with irinotecan cations [42]. Other groups studying salt variations in irinotecan gradient loading have also found TEA_8_SOS creates liposomes of superior stability when challenged *in vitro*, possibly due to sucrose octasulfate’s ability to promote “inter-fiber crosslinking” by interacting with multiple drug cations simultaneously [43]. These positive findings in the literature thus encouraged the pursuit of similar results for other drug cations such as NIRB and DOX.

To explore the potential utility of TEA_8_SOS as an effective trapping agent, it was important to determine whether NIRB and DOX could be co-loaded into TEA_8_SOS liposomes at the 10:1 molar ratio in a reproducible fashion. As shown in Figure 3, co-loading of NIRB and DOX into TEA_8_SOS aqueous core liposomes followed a linear relationship wherein incremental increases in NIRB relative to DOX resulted in linear incremental increases in NIRB:DOX encapsulated.

The linear regression model of this relationship (*y* = 1.10*x* – 0.178) was used to predict the drug loading parameters necessary to obtain our desired synergistic payload – i.e., that an addition of 9.24-fold molar excess of NIRB relative to DOX should yield liposomes loaded with near to the desired 10:1 NIRB:DOX molar ratio. This prediction was experimentally validated and led to the resulting NIRB-DOX-FA liposomes encapsulating 9.4±0.6:1 of NIRB:DOX, confirming TEA_8_SOS’s reproducible ability to co-entrap both drugs at high efficiencies.

It should be noted that pH gradient mediated active loading using sodium citrate (NIRB-DOX-CIT) as an intraliposomal entrapment agent was also attempted given its frequent use in the preparation of DOX containing liposomes [44]. Entrapment using sodium citrate, however, yielded NIRB and DOX liposomes with *in vitro* release characteristics that were less favorable than that of the current candidate, as shown in Figure S1. Specifically, co-encapsulation of NIRB and DOX into the core of sodium citrate liposomes resulted in rapid *in vitro* release. While NIRB-citrate liposomes maintained approximately 50% of their drug cargo after 120 hours of incubation (Figure S2), NIRB-DOX-CIT liposomes exhibited rapid release (~62% DOX and ~85% NIRB) at the 24-hour timepoint (Figure S1). We thus concluded that sodium citrate was an unsuitable entrapment agent for the co-loading of NIRB and DOX.

### 3.5 Cryogenic electron microscopy of NIRB-DOX liposomes reveal presence of an electron-dense drug core

Cryo-TEM images of NIRB-DOX-FA liposomes were taken to determine the morphology of both drugs at the liposomal core. As previously noted, a solid core is desirable to minimize drug release from FRα targeted liposomes while in systemic circulation. Cryo-TEM images of unloaded and dual loaded TEA_8_SOS liposomes presented in Figure 5b show that drug loading results in the formation of dark, electron-dense deposits at the liposomal core.

As shown in Figure 6, calculated mean gray value differences on the inside versus outside of NIRB-DOX-FA liposomes were found to be significantly larger than that of unloaded TEA_8_SOS core liposomes (p<0.0001), suggesting the presence of large, concentrated amounts of co-encapsulated drug within the internal volume of the liposomes. Since cryo-TEM imaging measures electron density, darker visible regions of the images represent areas of high encapsulated concentration of drug which, due to the work performed by others on DOX loaded liposomes, suggests the presence of drug precipitation which likely resulted in the observed payload stability [41].

Interestingly, per Figure S4, NIRB-DOX-FA exhibited a statistically significant increase in mean gray difference when compared to NIRB-DOX-CIT, suggesting a difference in sodium citrate’s ability to enable drug precipitation when compared to TEA_8_SOS. It should also be noted that internal media concentrations of anions for NIRB-DOX-CIT and NIRB-DOX-FA liposomes are 900 mEq and 650 mEq, respectively. As such, despite providing more available anions for electrostatic interactions with drug cations, sodium citrate was unable to facilitate drug precipitation relative to TEA_8_SOS, pointing towards sucrose octasulfate’s structural ability to induce drug precipitation which reduced the rate of drug release.

### 3.6 NIRB-DOX-FA liposomes exhibit sustained in vitro release profiles

*In vitro* release studies (Figure 4) in the presence of BSA (50 mg/mL in HBS) revealed that NIRB-DOX-FA liposomes exhibited no statistically significant amount of drug release over the duration of the study (comparing encapsulated drug levels at t=0 h vs t=120 h, p=0.9 and p=1.0 for NIRB and DOX, respectively). At the 120-hours timepoint, the encapsulated NIRB:DOX molar ratio remained close to the synergistic target.

As highlighted above, it is hypothesized that the observed high level of payload stability is due to the co-precipitation of both NIRB and DOX within the liposomal core. Precipitation has been shown to slow the release kinetics of drugs [40,43]. Had NIRB and DOX not formed intraliposomal drug precipitates, *in vitro* drug release would have likely followed a more rapid Fickian profile (i.e. due to drug release being governed by the concentration gradient across the liposome membrane) [45]. Thus, the hypothesis that NIRB-DOX-FA liposomes exhibit a solid core offers a more plausible explanation for their slow-release profile since drug release would additionally be governed by solid drug dissolution. Further characterization studies using techniques such as x-ray diffraction would be required to objectively confirm the state of the intraliposomal drug molecules.

Inherent stability of the dual-loaded formulation is ideal for active targeting to the characteristically overexpressed FRα to ensure controlled delivery of the identified synergistic ratio. Successful application of the active targeting strategy was similarly employed by Merrimack Pharmaceuticals Inc. for MM-302, a liposomal formulation of DOX actively targeted to the HER-2 receptor. As shown in the case of MM-302, active targeting leads to increased tumor cell uptake of the liposomes via receptor mediated endocytosis and improvement in the overall therapeutic effect [46,47].

## 4. Conclusion

At present, the above findings are the first (to our knowledge) to show synergistic effects for the combination of NIRB and DOX. Through our screening, we have found that the synergistic action of NIRB and DOX is both molar ratio and cell line dependent. NIRB and DOX, applied at a 10:1 relative molar ratio, exhibit consistent synergistic or additive effects for all tested HR deficient cell lines.

Perhaps the most important finding of this study was that the DOX and PARPi combination can lead to antagonism in cell lines with HR proficiency (i.e., intact BRCA2 function). In PEO4, the NIRB and DOX combination showed antagonistic effects across all applied molar drug ratios. These findings are in agreement with that of previous work done on the OLP and DOX combination by our group wherein mean CI values were found to be consistently higher in PEO4 versus PEO1 [28]. Our screening therefore suggests that combinations of DOX and PARPis in general may be inappropriate for patients with HR proficient tumors – highlighting the need to better understand the action of not just monotherapies but drug combinations on cancer cell biology.

Combined delivery of drugs using nanoparticles is one of the only ways to achieve ratiometric combinations at diseased sites *in vivo*. Formulation of an optimal dual encapsulated nanoparticle brings to the forefront an additional set of considerations. Details such as dual drug retention and release require meticulous optimization to obtain the ideal candidate formulation.

In this paper, we demonstrate the feasibility of developing a highly stable, nanoparticle containing a synergistic, 10:1 molar ratio of NIRB:DOX actively targeted with folate to promote receptor mediated endocytosis and thus precision delivery of the molar drug ratio of interest. The formulation was designed through considering distinct drug entrapment techniques rather than optimization of the lipid composition. To our knowledge, this is the first time NIRB and DOX have been successfully co-encapsulated into a single nanoparticle.

Caveats to this study include the small number of OC cell lines screened. Screening the NIRB and DOX combination in a larger panel of OC cell lines (including additional platinum resistant cell lines) could identify the combination’s performance in a more diverse setting which may better model disease heterogeneity. As shown in our results with PEO1 versus PEO4, drug combination performance can be altered noticeably with either the functional loss or gain of one single gene. Evaluation in diverse cell lines with key mutations is thus crucial to elucidate the clinical characteristics of potential good versus bad responders.

## 5. Materials and Methods

### 2.1 Materials

Hydrogenated soybean phosphatidylcholine (HSPC), N-(Methylpolyoxyethylene oxycarbonyl)-1,2-distearoyl-sn-glycero-3-phosphoethanolamine with a 2000 molecular weight PEG chain (PEG_2k_-DSPE) and a 5000 molecular weight PEG chain (PEG_5k_-DSPE) were purchased from NOF America Corporation (Kanagawa, Japan). Cholesterol was sourced from Corden Lipids (Plankstadt, Germany). 1,2-distearoyl-sn-glycero-3-phosphoethanolamine-N-[folate(polyethylene glycol)-5000] folate-PEG_5k_-DSPE) was purchased from Nanocs (New York, USA). DOX HCl and NIRB free base were purchased from Tongchuan Pharma (Wujiang City, Jiansu, China). Sodium sucrose octasulfate was purchased from Toronto Research Chemicals (North York, Canada). Human OC cell lines were obtained as follows: PEO1 and PEO4 (ECACC, Public Health England, Salisbury, UK) were purchased from Sigma-Aldrich; HEYA8 was obtained from M.D. Anderson Cancer Center (Houston, USA); OVCAR8 was obtained from the National Cancer Institute (Biological Testing Branch, NCI, MD, USA). All cell lines were authenticated using STR profiling by the Centre for Applied Genomics Genetic Analysis Facility (TCAG, Toronto).

### 2.2 In vitro combination cytotoxicity studies

A panel of four OC cell lines were selected for drug combination screening. Cells were seeded onto 96-well plates at the seeding densities listed in Table 1. Cells were left to adhere for 24 hours before undergoing a 72-hour treatment with NIRB and DOX both as monotherapies and in combination at varying molar ratios of NIRB relative to DOX. Cellular viability was then measured using the acid phosphatase (APH) assay as described previously [28]. Briefly, 2 mg/mL of phosphatase substrate (Sigma Aldrich, Oakville, ON, Canada) was added to 0.1M sodium acetate buffer, pH 5.5 with 0.1% Triton-X-100 and heated to 37°C. Drug containing media was removed followed by a wash with warm phosphate buffered saline pH 7.4 (PBS) prior to the addition of 100μL of warm phosphatase supplemented buffer. The plates were then incubated at 37°C for 1 hour followed by the addition of 10μL of 1M sodium hydroxide to stop the reaction. The UV absorbance was then read on a Cytation-5 plate reader at 405 nm (BioTek, Winooski, VT, USA). Fraction affected was calculated as follows:

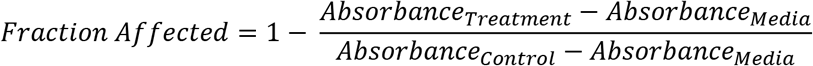

Total treatment concentration and the corresponding effects of NIRB and DOX both as monotherapies and varying relative molar ratios were inputted into CompuSyn software to obtain the combination index (CI) values of the treatments for fractions affected 0.5 and 0.75 (*f_a_* = 0.5 and *f_a_* = 0.75, respectively).

Developed by the Chou group, CompuSyn utilizes the median-effect equation to determine the Hill-type coefficient (m) and the dose required to induce a median cytotoxicity effect (D_m_) for both drugs singularly and in combination:

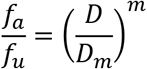

The median effect equation can be further manipulated to provide the basis of the median-effect plot:

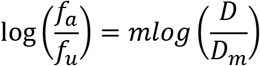

From the median effect plot, CompuSyn then calculates the doses of drug 1, drug 2, and the drugs in combination (i.e. (*D_x_*)_1_, (*D_x_*)_2_, and (*D_x_*)_1,2_, respectively) that elicit *x* fraction affected. These values are then inputted into following equation to calculate the CI value at varying fractions affected [48]:

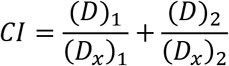

Where (*D*)_1_ and (*D*)_2_ are the doses of drugs 1 and 2 required to elicit fraction affected *x* when applied in combination. The resulting CI value quantitatively defines the combined dose molar ratio wherein CI values <1 are considered synergistic, equal to 1 additive, and >1 antagonistic. In practice, CI values representing synergy, additivity, and antagonism are herein conservatively defined as ≤0.9, between 0.9 and 1.1, and ≥1.1 respectively [28].

### 2.3 Evaluating FRα expression in panel of cell lines

FRα expression in the panel of OC cell lines was determined using flow cytometry. Briefly, the chosen panel of OC cell lines as well as the breast cancer cell line, MCF7 (used as a negative control) were plated onto 24-well plates overnight at a seeding density of 1.24 x10^5^ cells per well [52]. Cells were left to adhere for 24 hours before being scraped and collected. Treatment groups were incubated with an anti-FRα primary antibody (1:100, ab3361, Abcam, Toronto, ON, Canada) while control group cells were incubated with PBS at 4°C for 30 minutes. Cells were then pelleted via centrifugation, washed twice with ice cold PBS before incubation with the Alexa Fluor^®^ 488-labelled secondary antibody (1:2000, ab150113, Abcam, Toronto, ON, Canada) for another 30 minutes. After treatment with the secondary antibody, cells were pelleted and washed twice before resuspension in 500μL of cold PBS for analysis using flow cytometry (CytoFLEX 5, Beckman Coulter, Mississauga, ON, Canada) wherein mean fluorescence intensity of the gated cells (10,000 events) were collected and normalized to that of untreated cells (mean FITC treated divided by mean FITC untreated).

### 2.4 In vitro cellular uptake studies

Following analysis of FRα expression, the OVCAR8 and HEYA8 cell lines were selected as positive and negative controls, respectively. Liposomes passively loaded with the fluorescent probe calcein were prepared wherein HSPC, cholesterol, and DSPE-mPEG2000 were mixed at a 55:40:4.75 molar ratio and dissolved in ethanol at 70°C before direct hydration with 70mM calcein buffer (70mM calcein, 0.5mM EDTA, 0.5mM Tris in water, adjusted to pH 7.4 with 1M NaOH). Calcein liposomes were then post-inserted with 0.25% molar of either folate-PEG_5k_-DSPE or PEG_5k_-DSPE as the folate receptor targeted and folate receptor untargeted formulations, respectively. Liposomes were purified by dialysis against HEPES buffered saline (HBS) for 24 hours until dialysis buffer showed no visually discernable trace of fluorescent colour. The OVCAR8 and HEYA8 cell lines were maintained in folate-free RPMI media, for a minimum of 5 passages to ensure they were folate starved prior to commencing uptake studies. Folate starved OVCAR8 and HEYA8 cell lines were plated onto 24 well plates at a seeding density of 1.24 x10^5^ cells per well and allowed to adhere prior to treatment with either folate conjugated or unconjugated liposomes at a lipid concentration of 100μM phospholipid. Treated cells were then incubated at 37°C for 1 hour before triple wash with warm PBS. Washed cells were then scraped, collected, and analysed using flow cytometry as described above.

### 2.5 Formulation of unloaded liposomes

Unloaded liposomes were prepared using the following method. Briefly, HSPC, cholesterol, and PEG_2k_-DSPE were mixed at a 55:40:4.75 molar ratio and dissolved in ethanol at 70°C. Triethylamine sucrose octasulfate (TEA_8_SOS) buffer was prepared as described by Drummond et al. using ion exchange chromatography wherein the resultant free acid was titrated with neat triethylamine (TEA) to a pH of 5.7 prior to dilution to a final sulfate group concentration of 0.65M [53]. TEA_8_SOS buffer was then added to the dissolved lipids at 70°C to achieve a final phospholipid concentration of 100mM prior to being vortexed vigorously for two minutes to form multilamellar vesicles. The lipid mixture was then passed three times through double stacked, 0.2-micron pore size polycarbonate filters prior to extrusion through double stacked, 0.1-micron pore size polycarbonate filters (Whatman Inc., NJ, USA) 10 times using a 10mL Lipex Extruder (Vancouver, BC, Canada) at 65°C to make unilamellar liposomes. The resultant liposomes were then cooled on ice for 5 minutes before dialysis using a 50kDa MWCO regenerated cellulose dialysis bag (Repligen, California, USA) overnight in pH 7.4 HBS at 4°C.

### 2.6 Drug loading via active loading techniques

DOX and NIRB were dissolved in dimethyl sulfoxide (DMSO) and added to unloaded liposomes at a total concentration of 4% (v/v) DMSO and a 0.2 moles total drug:moles lipid (DL) ratio. Due to the limitations in allowable DMSO content, unloaded liposomes were diluted to an appropriate volume with HBS prior to addition of the drug solutions [54]. The resultant liposome and drug mixtures were then stirred at 65°C for 1 hour before being cooled on ice for 5 minutes. Loaded liposomes (NIRB-DOX) were then purified via size exclusion chromatography (SEC). SEC columns were packed using Sepharose CL-4B purchased from GE Healthcare Bio-Sciences (Mississauga, ON, Canada). Purified liposomes were then either diluted or concentrated by tangential flow to relevant concentrations for use.

### 2.7 Post-insertion of folate targeting ligand

Addition of the folate targeting ligand was achieved using the post-insertion technique developed by Uster et al. [55]. In brief, folate-PEG_5k_-DSPE was dissolved in HBS at 60°C at a 5 mg/mL concentration. The folate-PEG_5k_-DSPE solution was then added to the drug loaded liposomes such that the folate conjugated lipid accounted for 0.25% molar of the total lipid content. The liposomes were then stirred at 60°C for 1 hour. Resultant folate conjugated liposomes (NIRB-DOX-FA) were then purified by SEC to remove any free drug released during the post-insertion process. Unconjugated control liposomes were post-inserted with PEG_5k_-DSPE using the same technique as above.

### 2.8 Analysis of NIRB and DOX

The concentrations of NIRB and DOX in samples were quantified using both HPLC-UV and MS analysis. Briefly, NIRB and DOX were extracted from liposome samples by 10-fold dilution with methanol. The resultant samples were then centrifuged at 3000 RPM, 4°C for 15 minutes in glass centrifuge tubes to separate extracted drug from lipid. The supernatant was then assayed directly for HPLC-UV or diluted accordingly for MS detection.

HPLC-UV analysis of samples was performed using an Agilent 1260 infinity series LC (Agilent Technologies, Santa Clara, CA, USA). Chromatographic separation was achieved using an Agilent Eclipse XDB-C18 column (4.6×150mm, 5.0μm) at 25°C with a mobile phase composed of acetonitrile and methanol 50:50 (A) and 50mM ammonium acetate, pH 4 (B). The initial mobile phase was 40% A with a flow rate of 1.0 mL/min, which was gradually increased to 55% A over 4 minutes. Following a 4-minute equilibration, the composition was changed back to 40% A over a duration of 30 seconds. Mobile phase was then maintained for another 1.5 minutes until run completion.

Detection of both drugs was achieved using an Agilent 1260 Infinity II Diode Array Detector with detection of NIRB and DOX at 310 nm and 480 nm wavelengths, respectively.

HPLC-MS analysis was performed using an Agilent 1260 infinity series LC equipped with an Agilent EC-C18 column (2.1×50mm, 1.9μM) heated to 40°C. A gradient elution was applied using methanol (A) and water (B) both with 0.1% formic acid (v/v). The initial mobile phase was 75% A with a flow rate of 0.3 ml/min which was gradually decreased to 0% A over 4 minutes. This composition was maintained for another 6 minutes before it was rapidly changed back to 75% A where it was maintained for the remaining 3 minutes of the run.

Mass spectrometry detection of NIRB and DOX was achieved using a ThermoFisher Scientific TSQ Endura Triple Quadrupole Mass Spectrometer (Mississauga, ON, Canada). Analysis was performed on positive ion mode with optimal ion source settings as follows: spray voltage of 3500 V, sheath gas of 5 a.u., auxiliary gas of 2 a.u., and ion transfer tube temperature of 275°C. Selected precursor ions for NIRB were m/z 321.2 → 180.0, 205.0, 207.0, 232.0, 304.083 while precursor ions for DOX were m/z 544.2 → 320.9, 345.9, 361.0, 378.9, 396.9. Collision energies ranged from 19.0 V to 42.0 V for NIRB and 11.2 V to 41.2 V for DOX.

Encapsulation efficiencies of drug were calculated using the following equation:

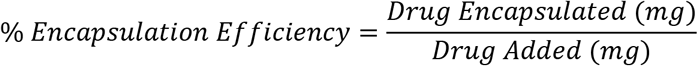

### 2.9 Lipid quantification

Quantification of HSPC and cholesterol was performed using an Agilent 1260 infinity series LC equipped with an Agilent evaporative light scattering detector (ELSD) operating at a 1.6 L/min gas flow with the evaporator and nebulizer heated to 80°C and 50°C, respectively. Lipids were extracted from liposome samples by direct dilution with methanol. Chromatographic separation of HSPC and cholesterol was achieved using an Eclipse XDB C18 column (150×4.6mm, 5μM) at 40°C with mobile phase composed of methanol (A) and water (B) both with 0.1% trifluoroacetic acid (v/v). The initial mobile phase was 90% A with a flow rate of 1.0 mL/min, which was gradually increased to 100% A over 4 minutes. Following an 11-minute equilibration, the composition was rapidly changed back to 90% A where it was maintained for the remaining 5 minutes of the run.

DL ratios were calculated using the following equation:

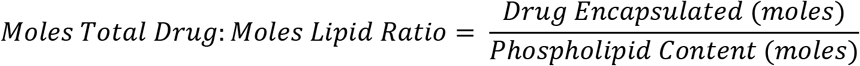

### 2.10 *In vitro* release of drugs

Evaluation of *in vitro* release of drugs from NIRB-DOX-FA was performed under sink conditions in 50 mg/mL of bovine serum albumin (BSA, heat shock fraction, pH 7, ≥98.0%, Sigma-Aldrich, Oakville, ON, Canada) over five days at 37°C. NIRB-DOX-FA liposomes were added to a BSA solution such that the final volume was 5 mL and the final concentration of NIRB and BSA were 1 mg/mL and 50 mg/mL, respectively. Samples of 100 μL were removed at pre-determined timepoints and purified by SEC to separate the liposomes from free drug. Sink conditions were maintained at all timepoints. NIRB and DOX were then extracted from the liposome fraction before analysis using HPLC-UV as described above.

### 2.11 Cryogenic transmission electron microscopy

Cryogenic transmission electron microscopy (cryo-TEM) imaging of NIRB-DOX-FA was performed at the University of Guelph Advanced Analytics Center. Liposome samples were diluted 50-fold with PBS and added to a Quantifoil Multi A holey carbon film (Quantifoil Micro Tools GmbH, Großlöbichau, Germany) deposited on a 300-mesh copper grid under humidity-controlled conditions at room temperature with excess sample removed prior to flash freezing with −183°C liquid ethane. Samples were then kept frozen by liquid nitrogen during the imaging process. Imaging was done using a FEI Tecnai G2 F20 microscope (FEI company, Hillsboro, OR, USA) equipped with a bottom mount Gatan 4k CCD camera (Gatan Inc., Warrendale, PA, USA). Images were taken in bright field mode at a 200 kV acceleration voltage.

Gray intensity of the imaged space both inside and outside the liposomes was measured using ImageJ Version 1.8.0 (NIH, USA) as mean gray area. Measurement of gray intensity was used to elucidate the physical state of the loaded drugs within the interior of the liposomes. The method used was adapted from the Cullis group [56]. Briefly, mean gray area expresses the gray intensity of all pixels in a selected area divided by the total number of pixels. Both internal and external areas of the imaged liposomes were sampled for mean gray area analysis. Internal mean gray values were then subtracted from external mean gray values to quantify the mean gray value difference across the bilayer. Approximately 50 liposomes were sampled from NIRB-DOX-FA images and compared to unloaded liposomes to elucidate the presence of concentrated NIRB-DOX deposits at the nanoparticle core.

### 2.12 Statistical Analysis

All statistical analyses were conducted using GraphPad Prism version 9.0. Mean gray value differences between loaded and unloaded liposomes were compared using unpaired t-tests. In addition, differences in FRα expression in the panel of OC cell lines were compared using one-way ANOVA and Tukey’s multiple comparisons test while differences in cellular uptake between folate conjugated and unconjugated liposomes in the same cell line were analyzed using paired t-tests.

## Supporting information

Supplementary Informaton

## 6. Acknowledgements

These studies were funded by the Department of Defense (Award number W81XWH-16-1-0388) and the Nanomedicines Innovation Network (NMIN). L.W. and L.A. acknowledge support from NSERC CREATE. L.W acknowledges scholarships from the Centre for Pharmaceutical Oncology (CPO). The authors acknowledge use of equipment in the CPO at the University of Toronto.

## 7. Author Contributions

**L.W.:** Conceptualization, Methodology, Validation, Formal Analysis, Investigation, Writing – Original Draft, Visualization; **J.E.:** Writing – Review and Editing, Project Administration; **L.A.:** Investigation, Writing – Review and Editing; **C.A.:** Writing – Review and Editing, Funding Acquisition, Supervision

## 8. Data Availability Statement

Data generated and analysed throughout the duration of this study are available upon request from the corresponding author.

## 9. Additional Information

The authors declare no competing interests.

## 10. Abbreviations

OC: ovarian cancer
APH: acid phosphatase
BSA: bovine serum albumin
CI: combination index
cryo-TEM: cryogenic transmission electron microscopy
DL: moles total drug:moles total lipid
DMSO: dimethyl sulfoxide
DOX: doxorubicin
dsDNA: double stranded DNA
ELSD: evaporative light scattering detector
EPR: enhanced permeability and retention
folate-PEG_5k_-DSPE: 1,2-distearoyl-sn-glycero-3-phosphoethanolamine-N-[folate(polyethylene glycol)-5000]
FRα: folate receptor alpha
HBS: HEPES buffered saline
HGSOC: high grade serous ovarian cancer
HSPC: hydrogenated soybean phosphatidylcholine
NIRB: niraparib
NIRB-DOX: untargeted niraparib and doxorubicin loaded liposomes
NIRB-DOX-CIT: sodium citrate core liposomes loaded with niraparib and doxorubicin
NIRB-DOX-FA: folate targeted niraparib and doxorubicin loaded liposomes
OLP: olaparib
PARPi: parp inhibitor
PBS: phosphate buffered saline
PEG_2k_-DSPE: N-(Methylpolyoxyethylene oxycarbonyl)-1,2-distearoyl-sn-glycero-3-phosphoethanolamine with a 2000 molecular weight PEG chain
PEG_5k_-DSPE: N-(Methylpolyoxyethylene oxycarbonyl)-1,2-distearoyl-sn-glycero-3-phosphoethanolamine with a 5000 molecular weight PEG chain
PLD: PEGylated liposomal doxorubicin
SEC: size exclusion chromatography
TEA: triethylamine
TEA_8_SOS: triethylammonium sucrose octasulfate

