## Supplementary Informaton for "Folate receptor targeted nanoparticles containing niraparib and doxorubicin for treatment of high grade serous ovarian cancer"

Supplementary Information

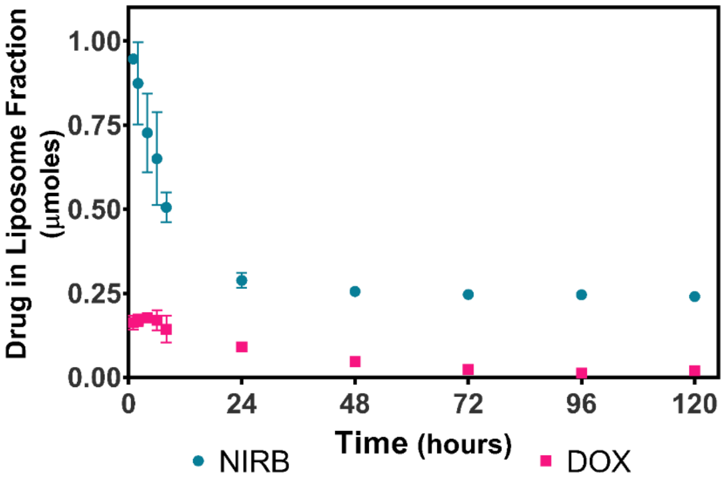


**Figure S1: *In vitro* release of NIRB and DOX from 300mM sodium citrate liposomes (NIRB-DOX-CIT) in BSA.** The y-axis shows the mean absolute value (µmoles) of NIRB and DOX remaining in the liposomes (±SD) within collected 100µL fractions when NIRB-DOX-CIT liposomes are subjected to biologically relevant (i.e., BSA; 50mg/mL) sink conditions over five days. Unlike sodium citrate core liposomes encapsulating only NIRB, dual-encapsulated NIRB-DOX-CIT liposomes exhibited rapid release of drug cargo within the first 24 hours (84.5±0.81% of NIRB and 61.9±6.49% of DOX at the 24-hour timepoint). The observed rapid release was unexpected considering the sustained release profiles of NIRB (shown) and DOX (per literature) liposomes when formulated individually using sodium citrate as an entrapment agent. It is hypothesized that while sodium citrate can adequately form stable ion complexes with either NIRB or DOX individually, the citrate anions are unsuitable for the entrapment of both NIRB and DOX simultaneously. Further investigation is required to elucidate the cause of this event. All experiments were completed as n=3.

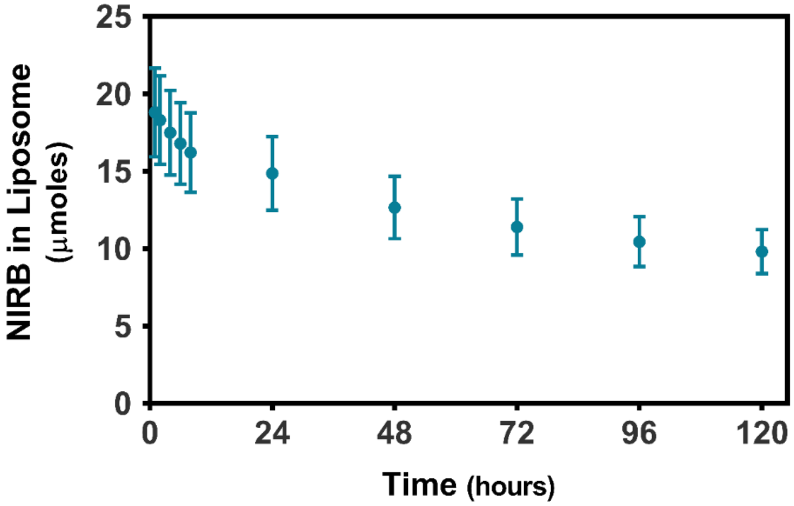


**Figure S2:** ***In vitro*** **release of NIRB from 300mM sodium citrate liposomes in BSA.** The y-axis shows the mean absolute value (μmoles) of NIRB remaining in all liposomes (±SD) when NIRB loaded sodium citrate core liposomes are subjected to biologically relevant (i.e., BSA; 50mg/mL) sink conditions over five days. Sodium citrate core liposomes encapsulating only NIRB exhibited sustained release over the five-day study (49.4±1.17% of drug was released at the 120-hour timepoint). All experiments were repeated as n=3.

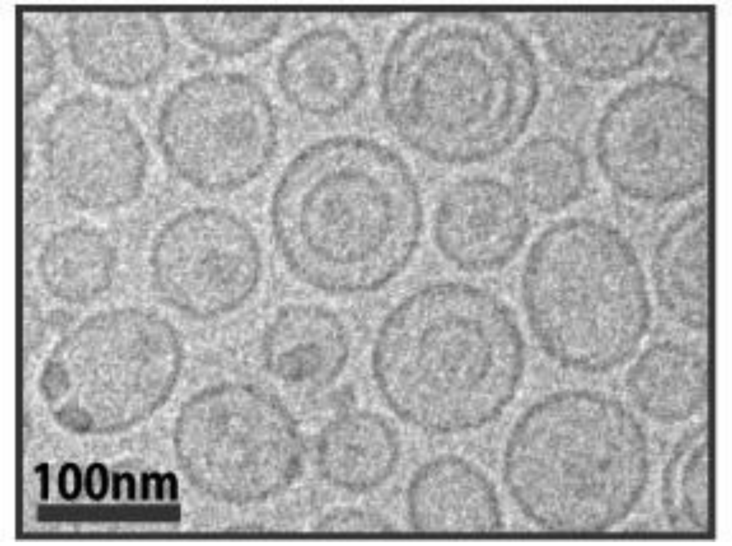


**Figure S3: Cryo-TEM images of folate conjugated, NIRB and DOX dual-encapsulated liposomes using sodium citrate as an entrapment agent (NIRB-DOX-CIT).** Note the lower levels of visually discernable intraliposomal dark areas, indicating a lack of electron dense deposits at the core. This is in direct contrast to NIRB-DOX-FA liposomes which use $TEA_8SOS$ as an entrapment agent, suggesting sodium citrate's inability to facilitate the formation of solid drug co-precipitates. These observed distinctions are plausible rationale for the stark differences in *in vitro* release profiles between the presented formulations.

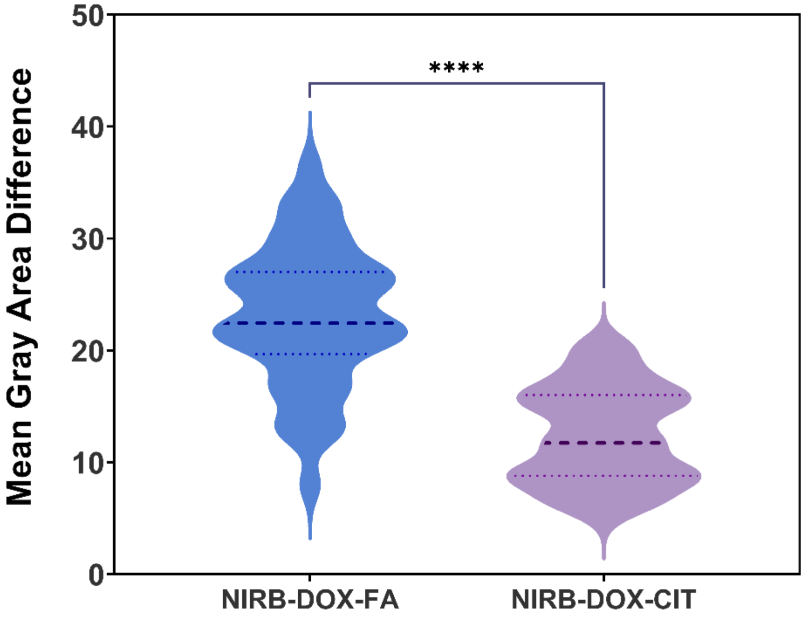


**Figure S4: Differences in mean gray area of the exterior versus interior of folate-targeted, NIRB and DOX dual-encapsulated liposomes using sodium citrate (NIRB-DOX-CIT) versus $TEA_8SOS$ (NIRB-DOX-FA) as an entrapment agent.** Mean gray area is defined as the average photo intensity divided by the number of pixels in a selected image area. Mean gray area was determined to quantify the darkened electron dense areas at the core of NIRB-DOX-CIT versus NIRB-DOX-FA liposomes. As such, statistically significant differences in mean gray area across the bilayer of NIRB-DOX-FA liposomes relative to NIRB-DOX-CIT liposomes, highlights the formation of electron dense drug deposits using $TEA_8SOS$ and not sodium citrate as an entrapment agent. Mean gray area of the liposomes was determined by ImageJ software for cryo-TEM images. Data is presented as differences in mean gray area (n=43, **** $p<0.0001$).
